# Understanding the Dynamics of Biomass Deconstruction b*y* the Cellulolytic Anaerobe *Clostridium thermocellum*

**DOI:** 10.1101/2024.11.19.624254

**Authors:** John M. Yarbrough, Dominik G. Stich, Neal N. Hengge, Qi Xu, Daehwan Chung, Samantha J. Ziegler, Shu Huang, Sarah Moraïs, Itzhak Mizrahi, Edward A. Bayer, Yannick J. Bomble

## Abstract

*Clostridium thermocellum* is one of the most efficient microorganisms for the deconstruction of cellulosic biomass. To achieve this high level of cellulolytic activity, *C. thermocellum* uses large multienzyme complexes known as cellulosomes to break down complex polysaccharides, notably cellulose, found in plant cell walls. The attachment of bacterial cells to the nearby substrate *via* the cellulosome has been hypothesized to be the reason for this high efficiency. The region lying between the cell and the substrate has shown great variation and dynamics that are affected by the growth stage of cells and the biomass used for growth. Here, we utilized both photoactivation localization microscopy (PALM) and stochastic optical reconstruction microscopy (STORM) in combination with Density-Based Spatial Clustering of Applications with Noise (DBSCAN) to study the distribution of *C. thermocellum* cellulosomes at different stages of growth when actively growing on soluble and insoluble substrates, providing a clearer picture of the dynamics of cellulosome populations at the enzyme microbe substrate interface. This research demonstrates the promising application of novel optical methodologies in tandem with targeted mutations within *C. thermocellum* to test the prevailing theories regarding the mechanisms of cellulosomes and their potential to shuttle onto the biomass for the attachment of *C. thermocellum* to improve biomass deconstruction.

## Introduction

Many microorganisms have evolved different mechanisms to efficiently deconstruct lignocellulosic biomass [1]. To date, *Clostridium thermocellum,* a thermophilic anaerobe, is the most efficient biomass degrader found in the biosphere with the capacity to solubilize more than 55% of the polysaccharide content of many key biomass feedstocks [2]. It has been hypothesized that this high propensity for biomass deconstruction is due to its complex and sophisticated cellulolytic machinery, which relies on supramolecular multienzyme complexes called cellulosomes [3–7]. These cellulosomes comprise non-enzymatic scaffolding proteins (scaffoldins) decorated with a wide range of complementary polysaccharide-degrading enzymes. In this cellulosomal system, complex formation is mediated by a strong interaction between the type I dockerin module of the enzymes and the complementary type I cohesin module found on the scaffoldins. The primary scaffoldin in *C. thermocellum,* CipA is composed of nine type I cohesin modules which can bind nine type 1 dockerin-bearing enzymes, a type II dockerin module, and a family 3 carbohydrate-binding module (CBM3a) (Figure 1) [8, 9]. These scaffoldins can themselves be assembled on secondary, type II cohesin-bearing scaffoldins, including the heptavalent OlpB, to form a supramolecular complex with up to 63 glycoside hydrolase enzymes. Some of these secondary scaffoldins are themselves attached to the microbial cell surface *via* a surface layer homology (SLH) domain, which appears to give these microbes and cellulosomes significant advantages over other microbes and free cellulases [9–13]. Moreover, the CBM3a allows the scaffoldin to selectively bind to cellulose. As a result of these varied interactions, *C. thermocellum* has been shown to adhere very strongly to the biomass substrate [14, 15].

**Figure 1.**
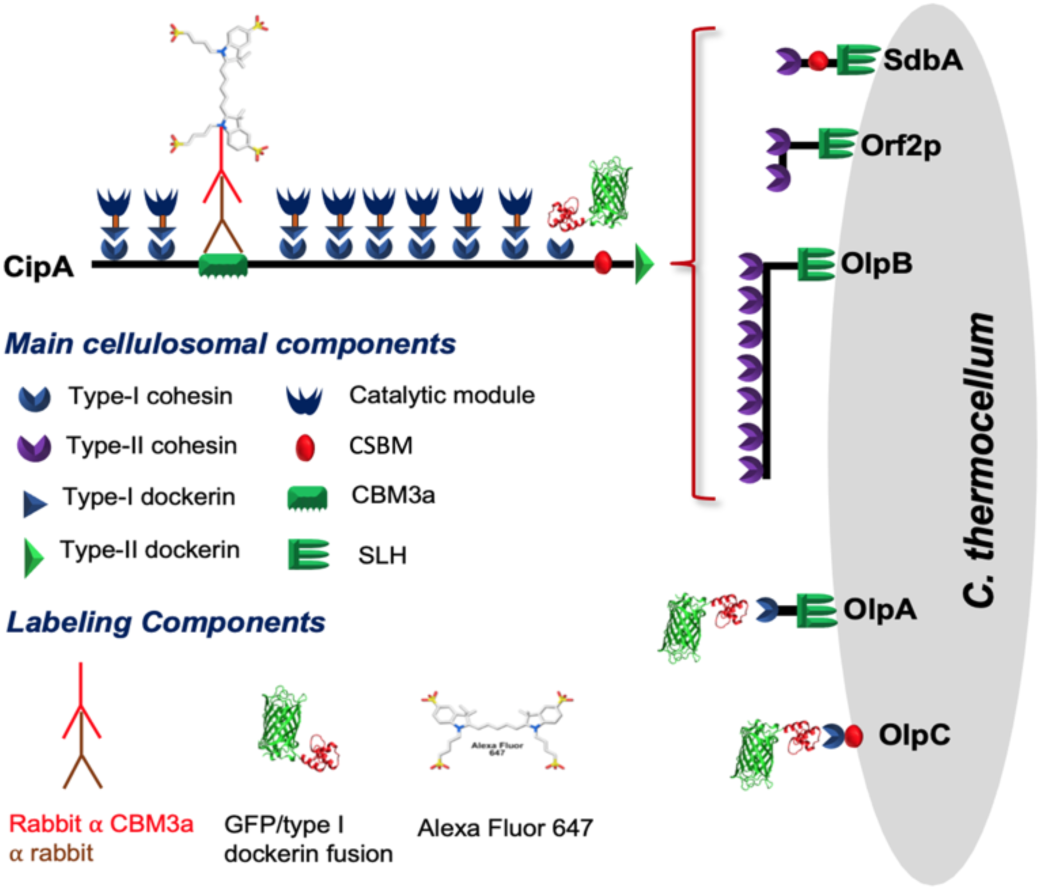
Main cellulosomal system of C. thermocellum, including cell-bound cellulosomes and components of our labeling strategy, using a combination of antibody-targeted chromophores specific to the family 3 carbohydrate-binding module (CBM3a) and a Green Fluorescent Protein-dockerin fusion protein (GFP-dockerin) fusion protein along with the Surface Layer Homology (SLH) domain and the Cell-Surface Binding Module (CSBM)

*C. thermocellum* has been extensively studied over the years to address many scientific questions regarding this elaborate cellulolytic system. Critical questions raised include: 1) What is the importance of the scaffoldins and the attachment of the cellulosome to the microbe? 2) How do cellulosomes and their composition change during growth, and are they released into the media?

Given the extensive literature regarding *C. thermocellum* and cellulosomes, we can now paint a clearer, albeit still incomplete, picture of the elaborate functioning of its cellulolytic machinery. For example, it has been shown that the bacterial cell walls are decorated with cellulosomes by Bayer and Lamed who were the first to show protuberance-like nodules on the outer surface of the bacterial cell walls using transmission electron microscopy (TEM) [14]. In the same work, they also showed that the conformation of cellulosomes changed when engaged on cellulose. Further, Bomble and coworkers were able to demonstrate, using molecular modeling, that even though dockerin-cohesin interactions seem to be similar for all cellulosomal glycoside hydrolases, the assembly process on the CipA scaffoldin was not random, but rather depended on the size and flexibility of these biomass degrading enzymes [16]. This notion garnered experimental support by evaluating the incorporation of different cellulosomal enzymes into a trivalent mini-scaffoldin [17]. More recently, Xu and coworkers took steps to develop several genetically modified strains of *C. thermocellum* to selectively delete the different scaffoldins that make up its cellulosome system, including those that tether cellulosomes to the bacterial cell wall [9]. Overall, these genetically modified strains were still able to degrade crystalline cellulose and biomass with the exception of the CipA-deleted strain which led to an almost complete inactivation of cellulolytic activity of the microbe [9]. Thus, the presence of the primary CipA scaffoldin is intrinsic to the observed efficient degradation of cellulosic biomass. Interestingly, some of these deletions also led to a change in the biomass deconstruction mechanism used by this microbe. The deconstruction mechanism of wild-type *C. thermocellum*, which proceeds via defibrillation of the biomass, can be attributed to its attachment to the biomass surface and, notably, the dynamics of the so-called Enzyme Microbe Substrate (EMS) interface located between the microbe and biomass surfaces. This reactive region has been hypothesized to be a key to the high cellulolytic activity of this microbe.

Several approaches can be used to study the EMS interface between *C. thermocellum* and biomass, such as scanning electron microscopy (SEM) or transmission electron microscopy (TEM) with immunolabeling. The latter, for example, was used by our group to characterize these interactions with limited success (data not shown). Both TEM and SEM can technically achieve the resolution needed to study the cellulosome, with resolutions around 0.2 nm [18, 19] and 2 nm [20, 21] respectively. However, the primary drawback of these techniques for this application is the extensive sample preparation needed, which can lead to long time lags between the main experiments and imaging of the sample. Furthermore, there is a lack of reliable statistics to analyze the level of immunolabeling due to difficulties in quantification. Optical microscopy can help alleviate the need for cumbersome sample preparation and improve quantitative analysis. However, until recently, the resolution of these techniques (greater than 250 nm in most cases) was restricted due to the diffraction barrier of light [22]. Nevertheless, within the last 10 years, a new field of optical microscopy has emerged that is capable of capturing the location of single fluorescent molecules with an optical resolution between 10 nm and 30 nm rendering them particularly well suited for more accurate localization of target molecules under near-native conditions [22–24] (See supplemental material).

Here, we utilized both photoactivation localization microscopy (PALM) [25] and stochastic optical reconstruction microscopy (STORM) [23, 26] to study the distribution of *C. thermocellum* cellulosomes at different stages of growth when actively growing on soluble and insoluble substrates, which provides a clearer picture of the dynamics of cellulosome populations at the EMS interface. We show in this work that 1) *C. thermocellum* bacteria grown on soluble substrates, exhibit large cellulosome clusters in the log phase that become smaller over the course of growth, 2) the concentration of cellulosomes in the EMS region, when bacteria are actively engaged on insoluble substrates, is far greater than previously expected, 3) bacteria actively bound to insoluble substrates increase the local concentration of cellulosomes at the contact points, and 4) bacteria that are not bound to the substrate (most likely after leaving the substrate) are depleted in surface-attached cellulosomes.

## Results

### I. Establishing the use of super high-resolution imaging of *C. thermocellum* on different substrates using multiple probes

Early studies have reported that the presence of cell-free cellulosomes on the surface of cellulose may be related to the detachment of the bacterium from the substrate [8, 9, 14, 15]. These studies also noted a reduction of cell-associated cellulosomes in the stationary phase of growth compared with the log phase. It was further proposed that detachment of the bacterial cells may be connected to a controlled release of the cells from the cellulose-bound cellulosome, which would continue to hydrolyze the substrate. Moreover, a recent study (Tatli et al, ref 30) that examined at nanoscale resolution single *C. thermocellum* cells in different growth phases using Cryo-ET, observed phenotypic cellulosomal heterogeneity mediated by soluble sugar concentration in the media, thus suggesting a division-of-labor strategy. We therefore sought to clarify further the status of cellulosomes with regards to the presence of both the primary scaffoldin and the cellulosomal enzymes on cells of *C. thermocellum* and on the substrate during the different phases of growth, using advanced optical microscopic approaches.

To enable the imaging of different components of the cellulolytic machinery of *C. thermocellum,* we used a primary antibody targeting the CBM3a of the CipA scaffoldin with a secondary antibody fused to Alexa Fluor 647 [26, 27], in combination with an engineered photoactivatable GFP (PA-GPF) [28], fused to a type-I dockerin that specifically targets type-I cohesins found on the CipA, OlpA, and OlpC scaffoldins, for STORM and PALM imaging, respectively (Figure 1). Figure 2 shows the fluorescence of these two probes in conjunction with *C. thermocellum* cells grown on cellobiose in the stationary phase. These super high-resolution optical images are reminiscent of the work conducted by Bayer and coworkers, wherein transmission electron micrographs of *C. thermocellum* showed similar protuberance-like structures [15]. Bayer and coworkers discovered these protuberances on *C. thermocellum* and reported that both their size and longitudinal arrangement varied as determined by either a selective cationic electron-dense probe or antibodies specific to the CipA scaffoldin [15]. Using STORM and PALM imaging, we were able to further validate these original findings, with minimal sample preparation and disruption of the bacterium, with a resolution of 35 nm. To date, these cellulosomal protuberances and their dynamics have not been fully characterized, especially on process-relevant feedstocks. These advanced optical microscopy techniques appear to be particularly well suited to the quantitative study of the population of cellulosomes and their dynamics over time during biomass deconstruction.

**Figure 2.**
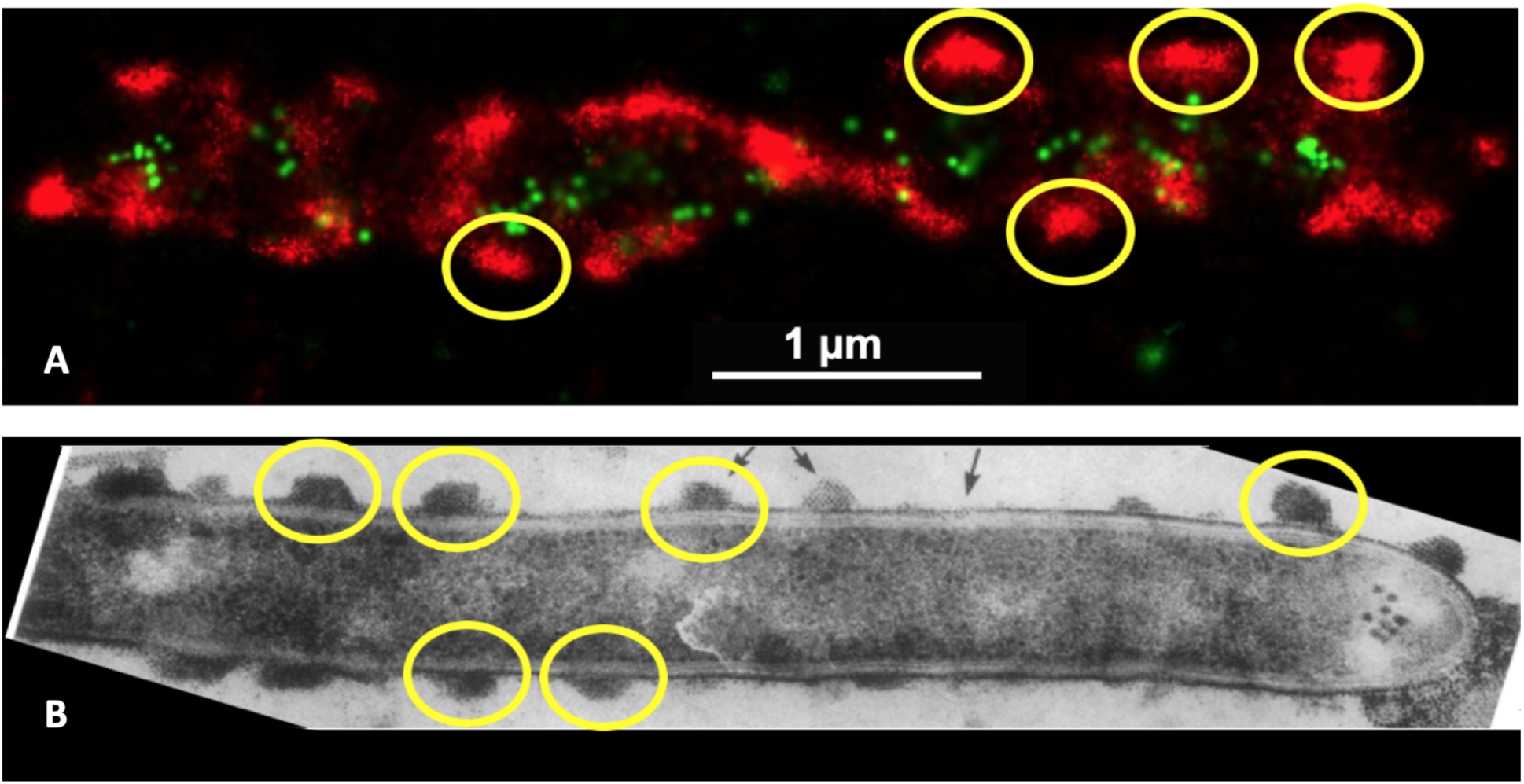
Distribution of cellulosomes on the surface of C. thermocellum. (A) Super high-resolution optical image of a C. thermocellum cell with the CBM3a located on the CipA scaffoldin tagged with Alexa Fluor 647 (red fluorescence) and a PA-GFP type-I dockerin fusion protein (green fluorescence) associated with type-I cohesins of either CipA, OlpA, or OlpC scaffoldins and (B) Adapted figure from Bayer et al. Ultrastructure of the cell surface cellulosome of Clostridium thermocellum and its interaction with cellulose showing a transmission electron micrograph of C. thermocellum, labeled with cationized ferritin [15]. In these images, some of the cellulosome protuberances are highlighted with yellow circles demonstrating that the surface of the bacterium is decorated with populated CipA scaffoldin

In this context, Figure 3 shows the distribution of cellulosomes (as indicated by the presence of CBM3a) on *C. thermocellum* in the log and stationary phases when grown on cellobiose. From these data, there is a distinct change in the pattern of the polycellulosomal protuberances which form highly decorated and interconnected protuberances on the bacterial cells in log phase. These new structures appear to be depleted when the microbe is in the stationary phase of growth. The fused GFP-dockerin probe also allows us to visualize the empty type-I cohesin positions on the primary scaffoldins CipA as well as those on scaffoldins OlpA and OlpC that are on to cell surface (Figure 1). Surprisingly, it seems that most empty type-I cohesins are located on the bacterial cell wall and that the CipA scaffoldins are almost fully saturated with enzymes.

**Figure 3.**
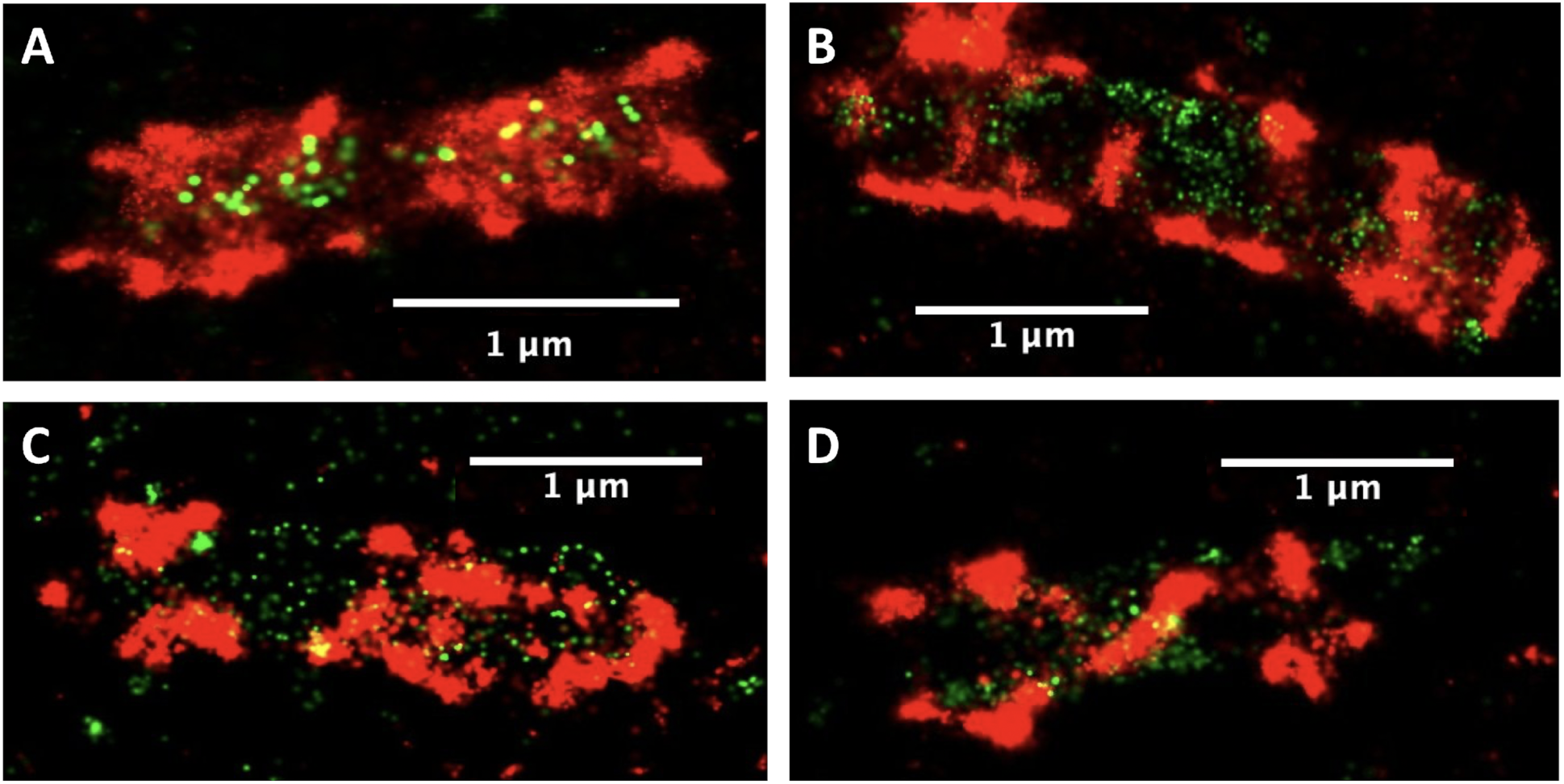
C. thermocellum grown in the presence of cellobiose in exponential growth phase (A, B) with an increase in the number of cellulosomes decorating the bacterial cell wall, in contrast to bacterial cells captured in the stationary phase (C, D) wherein the CipA scaffoldins are still present but with significantly reduced abundance. Red fluorescence denotes CBM3a (CipA scaffoldin)-associated label, whereas green fluorescence indicates unoccupied type I cohesin-related sites (see legend to Figure 2 and text for more detail).

To determine if the behavior of cellulosomes was different when *C. thermocellum* was grown on insoluble substrates, we grew *C. thermocellum* on microcrystalline cellulose (Avicel) particles and imaged the bacteria and substrate, 24 hours after inoculation, using the aforementioned probes. Figure 4 shows the fluorescence corresponding to the CipA-CBM3a antibody and the GFP/type-I dockerin fusion construct with the bacterial cells clearly attached to the Avicel particles. Most cells appear to have fewer cellulosome clusters than the log phase cells, grown on cellobiose, except at the EMS interface. In addition to the red and green fluorescence of the CBM3a and the scaffoldins, respectively, the yellow fluorescence is indicative of a colocalization of the green and red fluorescent molecules, signifying that there is a CBM3a in close proximity to an unoccupied type-I cohesin.

**Figure 4.**
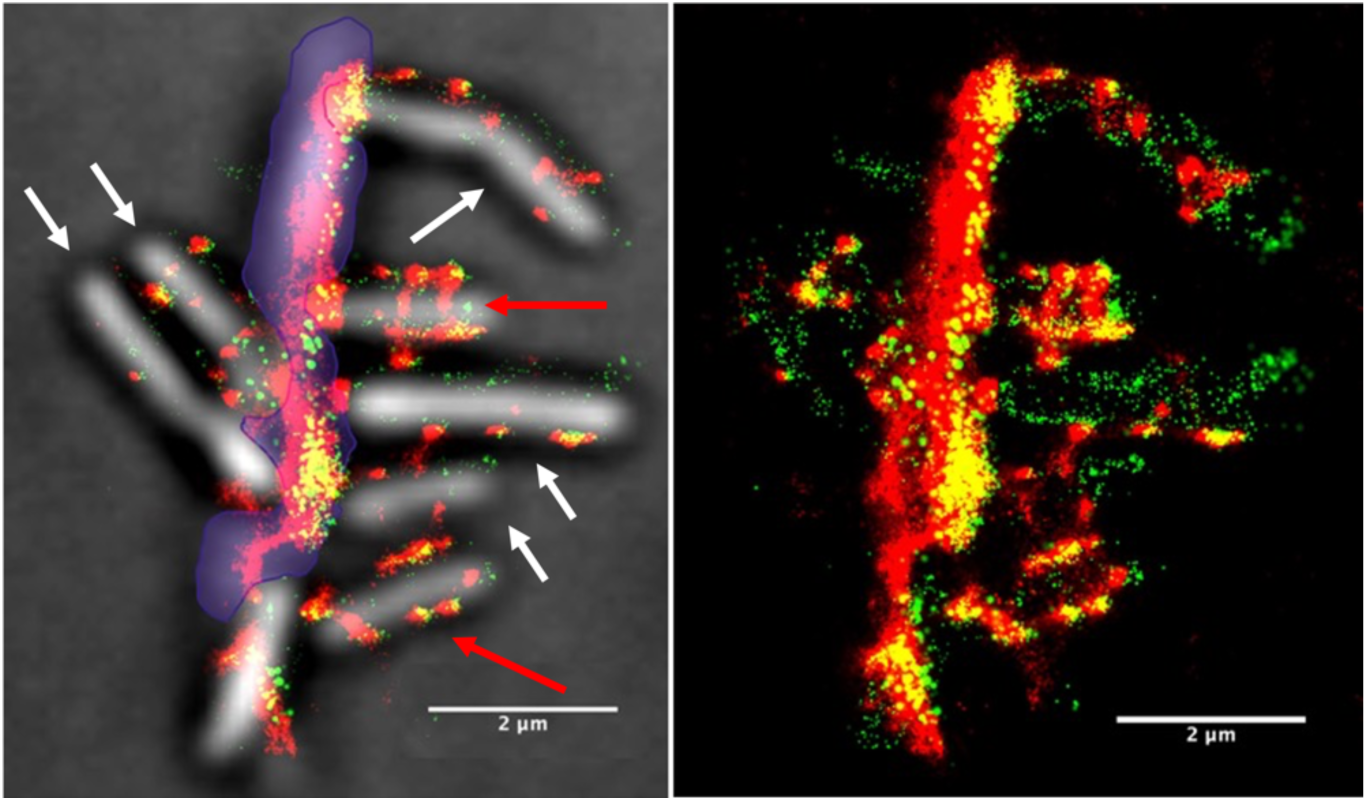
Images of C. thermocellum grown on Avicel captured in stationary phase showing. (A) the white light image superimposed with the fluorescence from the Alexa Fluor 647 (CBM3a) and the photoactivatable GFP/type-I dockerin fusion protein and (B) only the fluorescence from the Alexa Fluor 647 (CBM3a) and the fluorescence from the photoactivatable GFP/type I dockerin fusion. The Avicel particle is highlighted with purple shading in A. The increase in fluorescence within the EMS interface (area where the bacteria are connected to the Avicel) signifies that there is a high concentration of cellulosomes in that region.

The distribution of fluorescence on the Avicel particle indicates that there is a significant increase in CBM3a and unoccupied type-I cohesins at the EMS interface when the bacterium is attached to Avicel. The same observation can be made to some degree for cells binding to Avicel particles as well as on the rest of the particle itself (in the absence of bacterial cells). A similar increase in the fluorescence within this region was observed on other Avicel particles (see supplementary material), thus signifying that this high concentration of cellulosomes is not unique. Along with the increase in concentration of cellulosomes within the EMS interface, there is also a substantial amount of colocalization (yellow fluorescence) compared to cells grown on cellobiose, pointing to unoccupied, (*i.e.*, lacking type-I dockerin-bound enzyme) type-I cohesins (most likely from CipA) adjacent to CipA-CBM3a. Also, the fluorescence pattern on the bacteria seems to show two states, one in which the bacteria are decorated with CipA clusters (red arrows in Figure 4) and one in which the bacteria have few CipA clusters (white arrows) with lower fluorescence intensity, most likely indicating either that they are in different stages of growth or that they may have already relocated their cellulosomes to the contact points. These results agree with previous observations of phenotypic heterogeneity within *C. thermocellum* populations that were connected to a division-of-labor strategy [29].

In this section, we have shown that we can efficiently image different components of the *C. thermocellum* cellulolytic machinery using STORM and PALM on soluble and insoluble substrates with a resolution of close to 30 nm. These results clearly exemplify how powerful these techniques can be to answer many outstanding scientific questions about biomass deconstruction by *C. thermocellum*. However, current approaches to analyze super high-resolution images do not take advantage of the abundance of data that these techniques can provide. To remedy this problem, more quantitative approaches are needed to enable the high throughput analysis of tens to hundreds of bacterial cells.

### II. Quantitative analysis of STORM and PALM optical imaging

#### 1. A promising unsupervised machine learning approach

The main drawback of traditional optical microscopy is the inability to systematically quantify the information within the image. Unfortunately, with optical images, it is difficult to quantify the fluorescence from each bacterium, let alone analyze any significant portion of the bacterial cells. However, because a portion of the data generated from the super high-resolution images contains the x-y coordinates of the calculated gaussian for each fluorescent event, we can use this information to quantify the location and number of fluorescent events within a region of the image using unsupervised machine learning. This type of machine learning, also referred to as cluster analysis, allows us to define a certain epsilon radius (Eps) and set a minimum number of points, or in this case, the number of fluorescent molecules within the defined radius. The clustering algorithm chosen for this work was the density-based spatial clustering of applications with noise (DBSCAN), which is a propagative cluster detection method linking points that are closely packed together and detecting outliers using the user-defined epsilon radius and a minimum number of points or molecules [30]. Some of the main benefits of using DBSCAN for single molecule experiments are that DBSCAN can detect arbitrary shaped clusters, is quick, and is robust to outliers [30]. For the work presented here, we initially decided to set the epsilon radius to 75 nm for the CBM3a-tagged Alexa Fluor 647 molecules.

Figure 5 illustrates the DBSCAN algorithm with the green spheres representing the core points considered part of a cluster as they fulfill two criteria: 1) they are located within the epsilon and 2) they contain at least the minimum number of molecules (MNM). If a cluster is started, the algorithm will continue classifying each point and propagate the cluster, while each scanned point meets the given criteria for epsilon and the MNM. Once one of the criteria is not met, these points are classified as the border points (Border molecules, Orange Spheres in Figure 5). If a point does not meet the minimum number of points within the defined epsilon, then the point is classified as noise (purple spheres in Figure 5).

**Figure 5.**
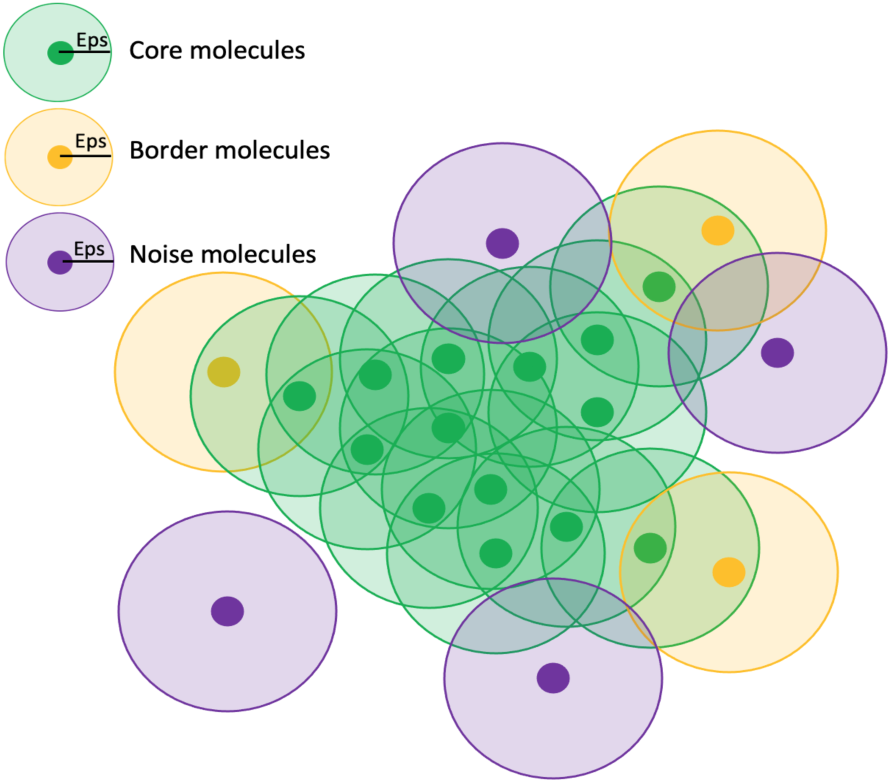
Graphical representation showing how the DBSCAN algorithm defines a cluster. The green spheres represent the core groups within the cluster, because they contain the minimum number of points within the defined eps radius. The orange spheres represent the border points which are still part of the cluster because they are within the defined epsilon but do not meet the minimum number of points criteria. The purple spheres indicate noise points and are not assigned to a cluster.

#### 2. *C. thermocellum* sheds cellulosomes over the course of growth on cellobiose

Figure 6 shows the distribution of cellulosomes (as indicated by the presence of CBM3a) on *C. thermocellum* in the log and stationary phases when grown on cellobiose. From these data, there is a distinct change in the pattern of the polycellulosomal protuberances which form highly decorated and interconnected protuberances on the bacterial cells in the log phase. These new structures appear to be depleted when the microbe is in the stationary phase of growth. The fused GFP-dockerin probe also allows us to visualize the empty type-I cohesin positions on the primary scaffoldins CipA as well as those on scaffoldins OlpA and OlpC that are tethered to the cell surface (Figure 1). Surprisingly, it seems that most empty type-I cohesins are located on the bacterial cell wall and that the CipA scaffoldins are almost fully saturated with enzymes. We utilized the DBSCAN cluster analysis with the number of clusters per bacterial cell versus the MNM per epsilon.

**Figure 6.**
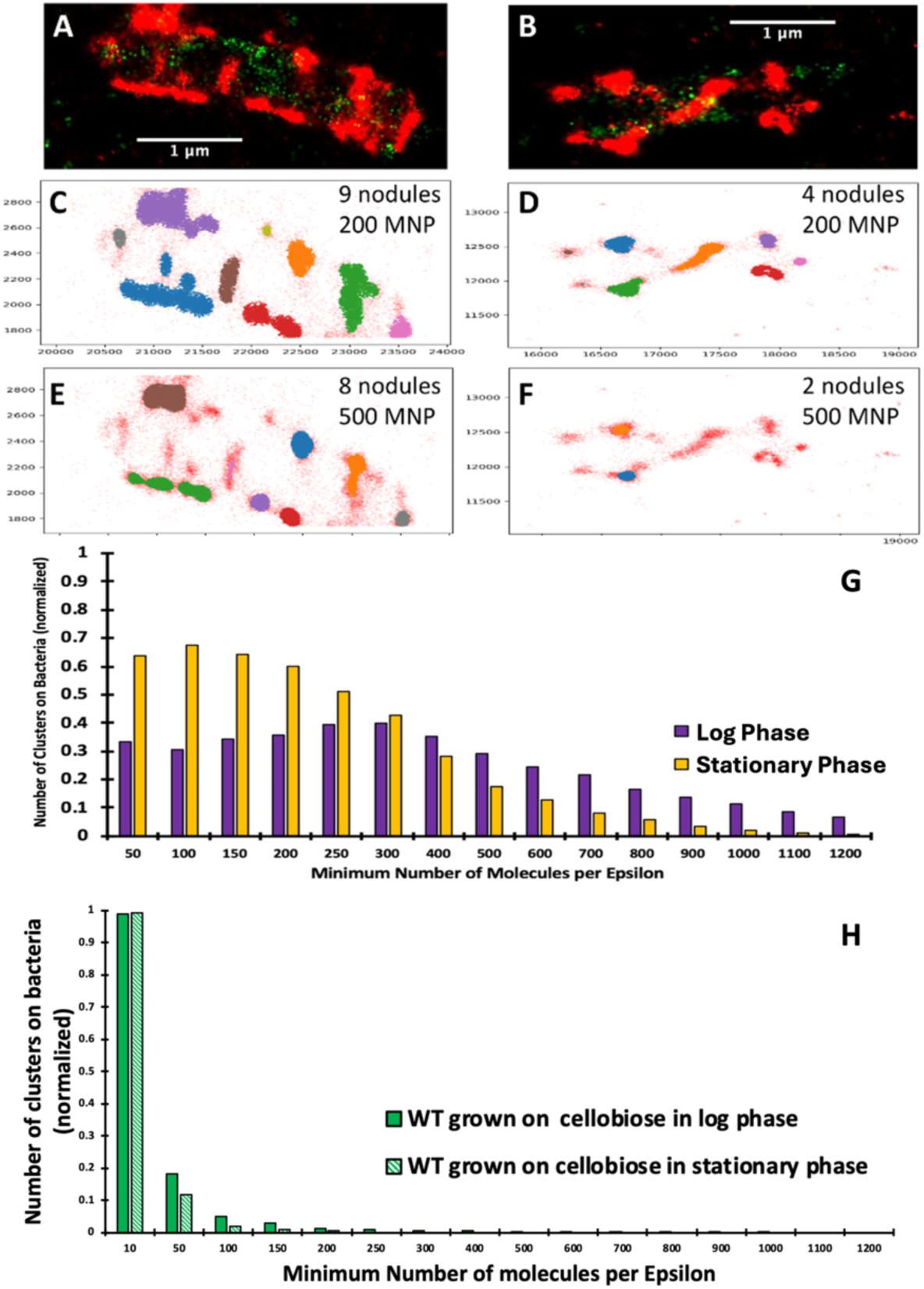
C. thermocellum grown in the presence of cellobiose in exponential growth phase. (A) with an increase in the number of cellulosomes decorating the bacterial cell wall, in contrast to bacterial cells captured in the stationary phase (B) wherein the CipA scaffoldins are still present but with significantly reduced abundance. Red fluorescence denotes CBM3a (CipA scaffoldin)-associated label, whereas green fluorescence indicates unoccupied type I cohesin-related sites (see legend to Figure 2 and text for more detail). DBSCAN results from C. thermocellum CBM3a-tagged cellulosomes, demonstrating the location and number of clusters for individual bacteria in the log phase (A) and stationary phase (B), with the number of detected clusters for 200 minimum number of molecules (MNM) per epsilon (C) and (D) and 500 MNM per epsilon (E) and (F). G. Cluster analysis of the cellulosomal protuberances on C. thermocellum in the log phase (purple) and stationary phase (yellow), normalized to 1. H. Cluster analysis of the unoccupied cohesins on C. thermocellum in the log phase (dark green) and stationary phase (light green), normalized to 1. The optical images are representative images from our data.

We also give a visual example of the number of clusters and their overall size on a specific bacterial cell in the log phase and in the stationary phases of growth. In this example, the number of cellulosome clusters found on the cell in stationary phase with a MNM per epsilon of 200 was four clusters (6E), and the number of clusters found for the bacterial cell in the log phase for 200 MNM per epsilon was nine (6C). When we increased the MNM per epsilon, in both cases, we saw a decrease in the number of detected nodules. For example, for 500 MNM per epsilon, two nodules were detected in the stationary phase, but eight nodules were detected in the log phase, thus illustrating that the nodules on the bacterial cells are larger in the log phase.

Altogether, we analyzed the normalized numbers of clusters collected from over 60 single bacterial cells in both the log and stationary phases of growth. Note that the x-axis represents the MNM per epsilon but does not give the relative size of the cluster, and the y-axis represents the normalized number of clusters per bacterial cell. The reasoning for normalizing the number of clusters per bacterial cell is to gain quantitative insight into how the density of molecules within the cellulosomal protuberances is affected during growth. During the log phase, there is an increase in the density of molecules compared to that of bacterial cells in the stationary phase. In the stationary phase, the number of identified clusters decreases as the density of molecules increases starting from 100 molecules per epsilon, whereas the maximum number of clusters per bacterial cell in the log phase is associated with 300 molecules per epsilon. This analysis was also used to determine potential clustering of unoccupied type-I cohesins on microbial cells (Figure 6H). The fluorescence corresponding to unoccupied cohesins shows limited clustering overall with only one cluster with 10 molecules per epsilon. Most importantly, the clustering of unoccupied cohesins on the *C. thermocellum* bacterial wall does not seem to follow the same pattern as that of the CipA-CBM3a, as the distribution in both the log phase (from 67 single bacteria cells) and the stationary phase (from 70 single bacteria cells) remains the same in both stages of growth. The signal for the unoccupied type-I cohesins and these analyses indicate that they are fairly dispersed on the surface and not co-localized with cellulosomes, which indicates either a low abundance of unoccupied type-I cohesins or a low abundance of type-I cohesins on the surface of these bacteria.

#### 3. *C. thermocellum* relocates cellulosomes over the course of growth when attached to insoluble substrate

To study the distribution of cellulosomes on bacterial cells and substrate during biomass deconstruction, we subjected *C. thermocellum* cells grown on Avicel (representative of an insoluble substrate) to the same imaging and analysis procedure. The bacterial cells were again captured during two stages of growth, log and stationery. Due to the added complexity of an insoluble substrate, each image set was inspected, and a python algorithm was used to select either individual bacterial cells, multiple bacterial cells (if the cells were too close in proximity to be captured individually), Avicel particles, or Avicel particles and bacteria together (See the Methods section for more details on this procedure).

Figure 7 shows a collection of images corresponding to *C. thermocellum* cells actively degrading Avicel during logarithmic growth. The Avicel particle in this figure is covered with multiple bacteria and identified as the underlying speckled red fluorescence with less intensity than the bacteria themselves (Fig 7C). This analysis shows the extensive network of CipA scaffoldins (cellulosomes) on both the bacterial cell wall and the Avicel particle (Fig 7D), whereas Figure 7E shows all unoccupied type-I cohesins on either the CipA scaffoldin or the bacteria (OlpA or OlpC scaffoldins). From this image, the complexity of the CipA scaffoldin network on both the surface of the bacteria and on the surface of the Avicel particle is quite visible and makes it rather difficult to draw any conclusion by simply comparing one section of the image to another. Therefore, these data, along with 15 other Avicel particles and 75 individual bacteria cells, were analyzed using the DBSCAN clustering algorithm. The clustering algorithm ran through an array of MNM per epsilon from 10 to 1200 at a fixed Epsilon radius set at 75 nm.

**Figure 7.**
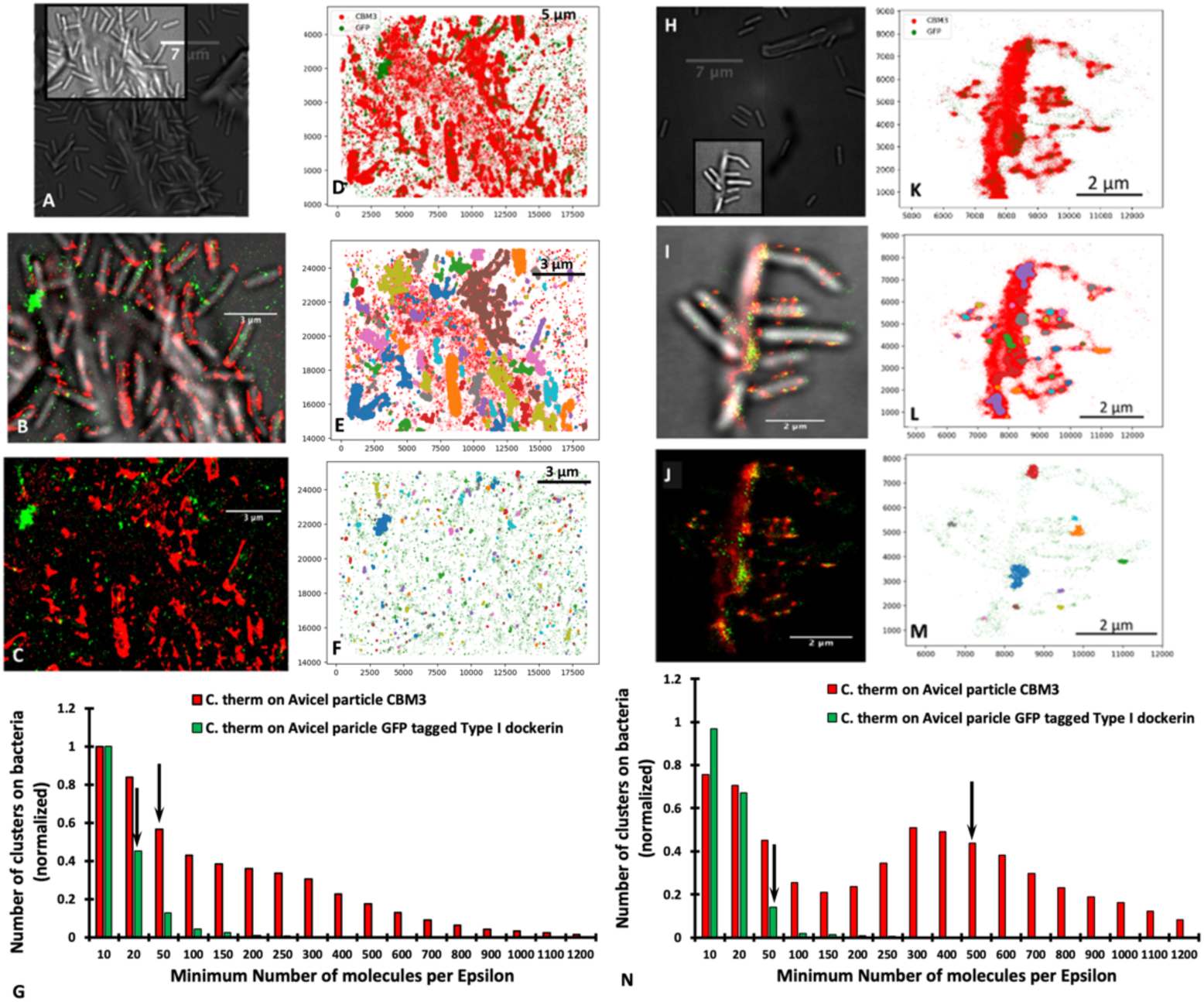
DBSCAN cluster plot of C. thermocellum grown on Avicel in the log phase. (A-G) and the stationary phase (H-N), showing (A,H) the white light image of the area analyzed by the clustering algorithm, (B,I) the overlay of the white-light and fluorescence image, (C,J) the fluorescence image alone, (D,K) the combination of the x-y positions for both the Alexa Fluor 647 and the photoactivatable GFP (individual plots for red and green fluorescence can be found in supplemental figure X), (E,L) the clusters identified for the Alexa Fluor 647-tagged CBM3a with an epsilon of 75 nm and a minimum number of molecules (MNM) set to 40 (E) or 500 (L), (F,M) the clusters identified for the photoactivatable GFP fusion protein with an epsilon of 125 nm and a minimum number of molecules of 20 (F) or 50 (M), and (G,N) the number of clusters vs the minimum number of points for CBM3a (red) and for GFP-dockerin (green). The arrows within the bar graph represent the minimum number of points chosen for the visual representation in (E, L – red) and (F, M – green).

The results of this analysis with the MNM set to 40 for the CBM3a (Figure 7D) show a total of 205 clusters of CipA-CBM3a identified, in which most of the clusters are located on the bacterial cell wall, thus demonstrating that there are high concentrations of CipA scaffoldins on the surface of the bacteria. It is very surprising to have more than 40 CipA scaffoldins packed in an area of ∼ 0.018 square microns, meaning that cellulosomes can be very tightly packed on the surface of the bacterium. For an MNM set to 500, there is a total of 127 clusters, again suggesting that cellulosomes can be very tightly packed within a small area, and these clusters are still mainly found on the surface of the bacterial cell wall. The cluster analysis for the PA-GFP tagged to the Type I dockerin with the MNM set to 20 with an epsilon radius of 125nm, led to the identification of 263 clusters. PA-GFPs are associated with either an unoccupied OlpA/OlpC site located on the bacterial cell wall or an unoccupied type-I cohesin located on the cellulosome, and for this data set, the clustering algorithm identified a total of 85 clusters that were primarily located on the bacterial cell wall. Figure 5G shows the distribution of the normalized number of clusters versus the MNM/cluster. Here we can see that for the PA-GFP tagged to the Type I dockerin, the distribution appears to be exponential or geometric, whereas the decrease in the distribution of CBM3a appears to be more attenuated. At this stage of growth, there appears to be no significant amount of CipA scaffoldins on the Avicel surface, compared to the bacterial surface, as indicated in Figure 5E. These samples collected midway through the log phase suggest that the majority of the CipA scaffoldins are attached to the bacterial cell, similar to that shown for the cellobiose-grown *C. thermocellum* during the log phase.

When analyzing the stationary phase when attached to Avicel, we found a significant amount of colocalization of the CipA-CBM3a and the type-I cohesin, which is primarily located on the Avicel particle and in particular at the point where the bacterium is attached to the Avicel particle (Figure 7H-J). Additionally, the Avicel particle seems saturated with cell-free cellulosomes at this stage. Figures 7L and 5M highlight clusters from the CipA scaffoldin molecules with the epsilon set to 75 nm with the MNM per epsilon set to 500 and clusters from the PA-GFP tagged to the Type I dockerin with the epsilon set to 125 nm with a MNM set to 50, respectively. This clustering analysis demonstrates two concepts, 1) there is a higher concentration of CipA scaffoldin molecules within 75 nm on the Avicel in comparison to the bacteria (as shown with the shaded region) and 2) the Type I dockerin clustering is primarily located on the Avicel particle. The analysis of these bacteria, shown in Figure 7N, reveals the distribution of the number of clusters on the bacteria vs the MNM per Epsilon, which appears to be slightly left skewed (longer on the left side of its peak than on its right). These results suggest that during the stationary phase, there is a higher concentration of CipA scaffoldin molecules that are tightly packed within the defined epsilon on the surface of the Avicel substrate. This is different compared to the concentration of CipA scaffoldins in the log phase. This demonstrates that during the life cycle of the bacterium, cellulosomes are gradually detached from the bacterial cell wall and adhere to the surface of the Avicel. This is exemplified by the center of the distribution being shifted toward a higher MNM per Epsilon and fewer overall numbers of clusters.

#### 4. *C. thermocellum* retains more cellulosomes during log phase when detached from insoluble substrate as compared to bacteria attached to substrate

The analysis of substrate-free bacteria revealed a remarkable distribution of CipA scaffoldins. The cellulosomes appear to cover the surface of the bacterial cell. Using our clustering algorithm, we can identify 14 clusters for the CipA-CBM3a containing at least 200 fluorescent molecules per Epsilon (Figure 7B) but only eight clusters for the PA-GFP containing at least 10 fluorescent molecules per Epsilon (Figure 7C). Figure 7D shows the normalized distribution of clusters for both the CBM3a (red) and the PA-GFP (green), averaged over 69 bacteria cells. The PA-GFP shows a very similar exponential or geometric distribution to that in Figure 5G, demonstrating that the combination of unoccupied OlpA/OlpC/CipA sites is similar for detached and attached bacteria. However, it appears that the distribution of the CipA-CBM3a clusters for the detached bacteria is quite different from the ensemble distribution shown in Figure 5G, which would indicate more of a mixed population in the culture. In the stationary phase, our analysis of detached bacterial cells revealed a different pattern than that observed for log-phase cells. The overall intensity and distribution pattern are significantly different and show a loss in the number of cellulosomes attached to the surface of stationary-phase bacteria (Figure 7E). Figures 7F and G show representative distributions for an MNM of 500 for CipA-CBM3a and 50 for the PA-GFP with 17 clusters detected for CipA-CBM3a and only seven clusters detected for PA-GFP (Figure 7H).

Considering the above data, we can now systematically compare the differences in distribution between the cellulosome and the unoccupied cohesins during different stages of growth for substrate-bound and substrate-detached bacterial cells. Figure 9A shows a clear shift in the distribution of cellulosomes on substrate-bound cells where significantly larger clusters are observed in the stationary phase. As illustrated in Figure 7I, most of these large clusters are found at the EMS interface between the microbes and the biomass, accompanied by a large increase in fluorescence. Figure 9B, however, shows a different phenomenon for detached bacterial cells with cells that seem depleted in cellulosomes that are in smaller clusters. The comparison of bound and detached log-phase bacterial cells shows that they have a very similar distribution of cellulosomes (Figure 9C). These results are illustrated in Figure 5B wherein microbes are bound to the biomass but do not seem to have relocated their cellulosomes to the contact point with biomass as is the case in the stationary phase. However, the difference between bound and free cells in the stationary phase is striking with very large clusters on bound cells but much smaller cellulosome clusters on free cells (Figure 9D). In contrast, the distribution of the unoccupied type-I cohesins appears to be unchanged under all conditions on bound and detached cells as shown in Figure 9E-H, indicating that the type-I cohesin vacancies do not change during the different stages of growth. This is an important result, as it has been hypothesized that some of the smaller scaffoldins (i.e., OlpA and OlpC) could serve as a shuttle to populate cellulosomes [31]. These results would seem to contradict this hypothesis as the number of unoccupied cohesins remains constant throughout growth.

**Figure 8.**
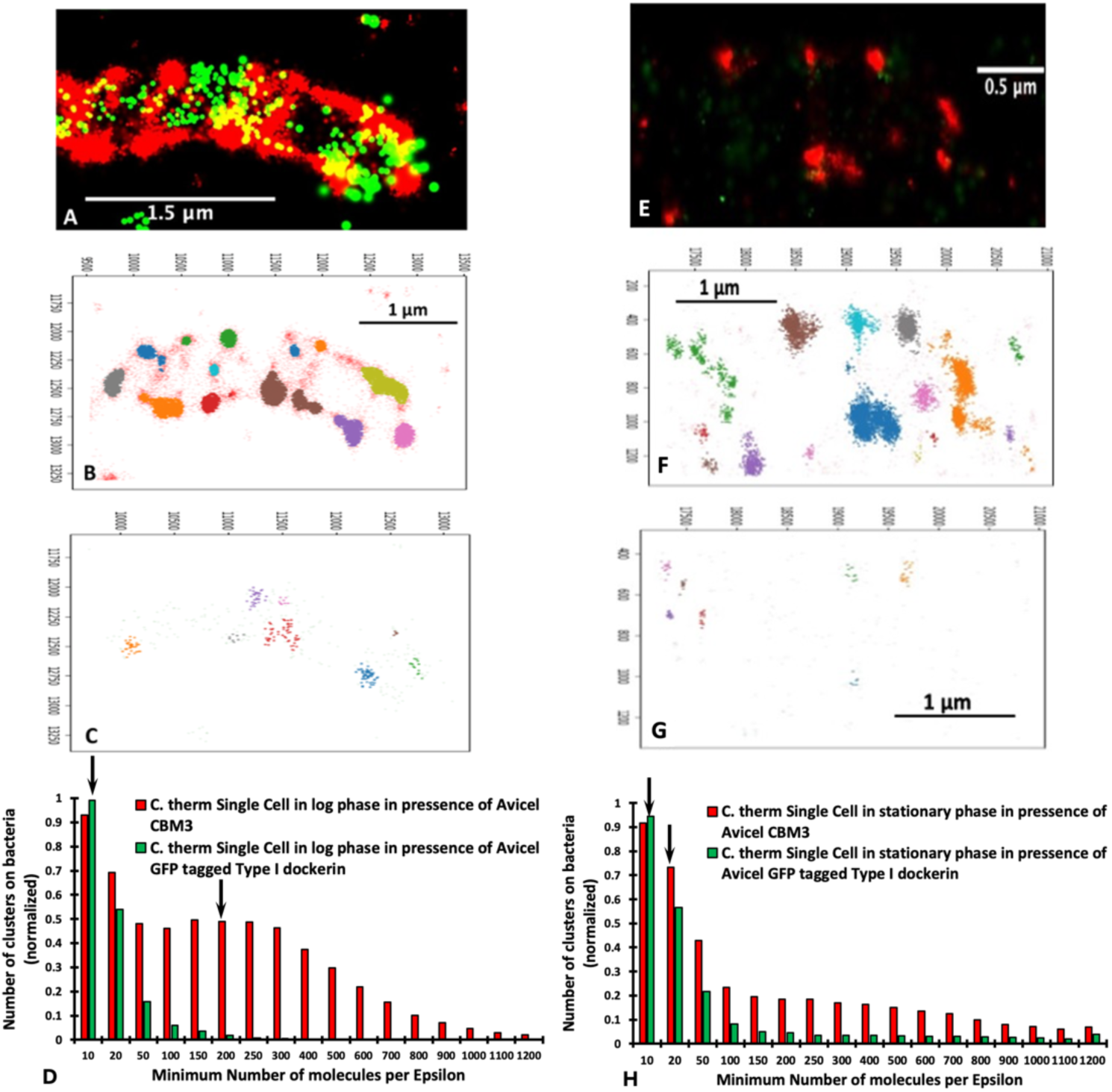
DBSCAN results from a detached C. thermocellum cell, grown on Avicel during the log. (A-D) and stationary (E-H) phases of growth, showing (A,E) the original fluorescence image, (B,F) the clusters identified for the Alexa Fluor 647-tagged CBM3a with an epsilon of 75 nm and an MNM set to 200 (B) or 20 (F), (C,G) the clusters identified for the fluorescent molecule of the photoactivated-GFP fusion protein with an epsilon of 125 nm and an MNM of 10 and (D) the number of clusters vs the MNM for the CBM3a (red bars) and for the PA-GFP (green bars). The arrows within the bar graph represent the MNMs chosen for the visual representation in (B,F) and (C,G) for a detached bacterial cell.

**Figure 9.**
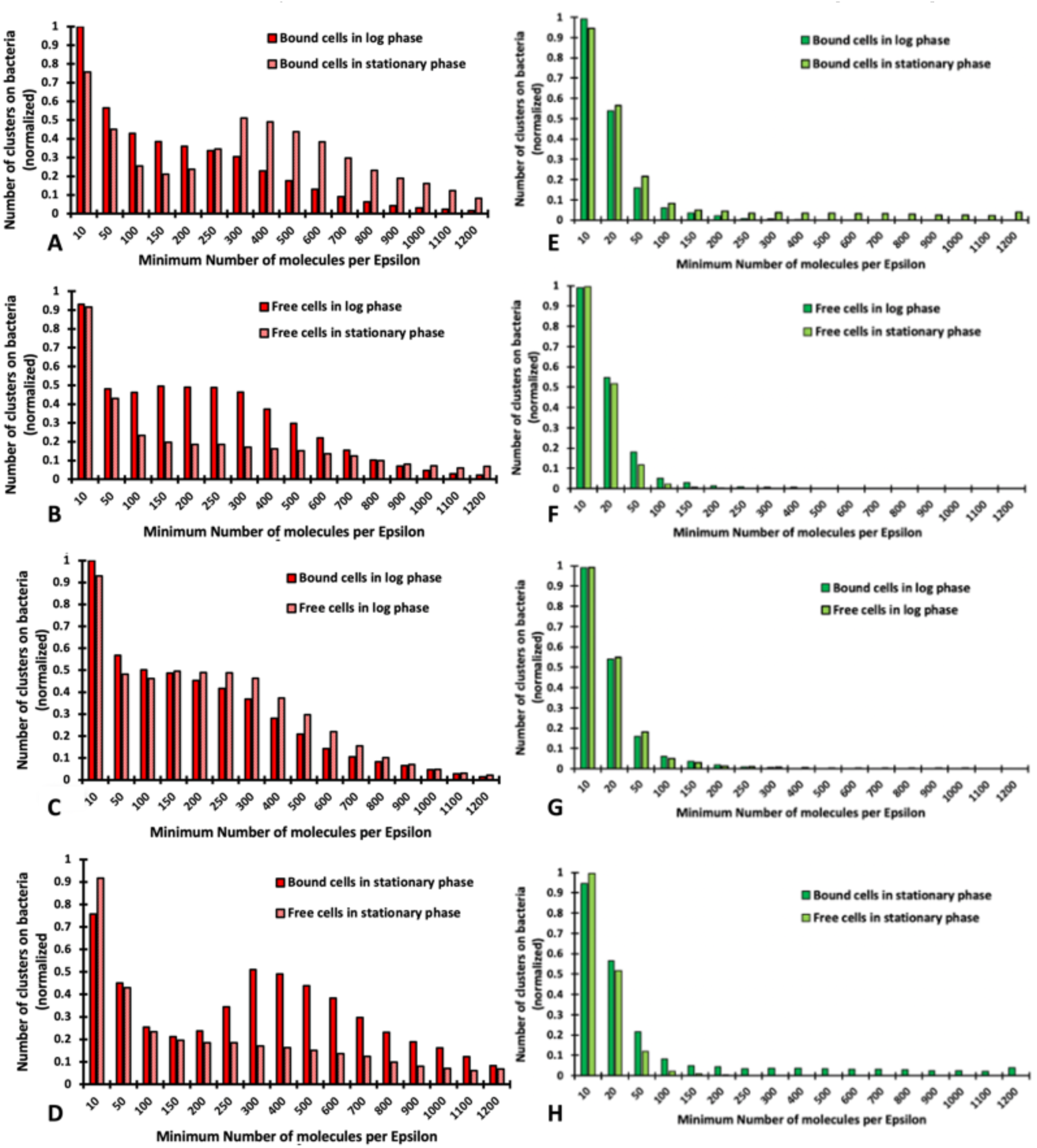
DBSCAN results focusing on the distribution of cellulosomes for C. thermocellum cells grown on Avicel, probing the effect of the same type of cells (bound or detached) being in log or stationary phase (A and B, respectively for bound or detached) and probing the effect of cells being bound or detached in each stage of growth [log phase (C) and stationary phase (D)]. Similar results focusing on the distribution of unoccupied type-1 dockerin are shown for C. thermocellum cells grown on Avicel, probing the effect of the same type of cells (bound or detached) being in log or stationary phase (E and F, respectively for bound or detached) and probing the effect of cells being bound or detached in each stage of growth (log phase (G) and stationary phase (H). These analyses show little differences in the distribution of unoccupied type-1 dockerins on the C. thermocellum cells in the different growth stages.

## Discussion

Biomass deconstruction by thermophilic microbes is a complex and heterogeneous process. This process is even more challenging to understand when considering cellulosomal microbes that primarily use an intricate multienzyme complex to deconstruct cellulosic substrates instead of free enzymes.

Over the years, considerable progress has been made toward understanding and exploiting cellulosomal systems to enable efficient deconstruction of cellulosic biomass. In early work to identify the *C. thermocellum* cellulosome, Bayer and Lamed observed nodulous protuberances on the surface of the microbe using transmission electron microscopy and immunolabeling [32]. Using super high-resolution techniques with sample preparation that more closely preserves native conditions, we observed similar protuberances in the present work (Figure 2). However, the distribution of these protuberances was different in cells grown to the log and stationary phases, when the microbe was actively engaged in degradation of cellulosic biomass. This finding agrees with a recent study that revealed a link between the stages of growth and the concentration of sugars and depletion of cellulosomes from the cell surface of cells grown on an insoluble cellulose substrate. Similar to previous work by tal] et al [29] we demonstrated herein that the microbes shed cellulosomes over the course of growth. We made the same observation when *C. thermocellum* was grown on both soluble and insoluble substrates, which seems to indicate that the loss of cellulosome is not triggered by their attachment to insoluble substrates. It is important to note that in the present study these conclusions were drawn from average fluorescence values, corresponding to cellullosomes from tens of bacteria grown on different substrates, stages of growth, detached or attached to biomass. Even though it appears that cells shed cellulosomes during growth on either soluble and insoluble substrates, it is clear from Figures 4, 7 and 9 that these cellulosomes are targeted to the biomass at the contact points in a concerted manner when substrate is present. Indeed, log-phase bound and detached cells appear to exhibit the same distribution profiles for the number of molecules per cellulosome cluster. However, in the stationary phase, detached bacterial cells bear much smaller cellulosome clusters, whereas bound cells exhibit much larger cellulosome clusters (Figure 9). We showed that stationary-phase bound cells in contact with the cellulosic biomass seem to reposition and concentrate the cellulosomes at the contact points or EMS interface (Figure 4). These results are consistent with early TEM observations [3, 15] whereby the exocellular protuberances of cellulose-bound cells were reported to undergo dramatic conformational change to form contact corridors where the cellulosomes are concentrated on the surface of the cellulosic substrate. Additionally, there seems to be increased colocalization of empty type-I cohesin and cellulosomes at these contact points (Figure 4), indicating that some of the dockerin-bearing enzyme molecules occupying these cohesin-bearing scaffoldins were released to interact directly with the substrate. The unoccupied type-I cohesin can be found on different protein scaffoldins, e.g., OlpA and OlpC, attached to the microbial cell wall, but also on the primary scaffoldin, CipA, either detached or initially bound to the bacterial cell wall via the secondary scaffoldin OlpB (Figure 1). In Figure 4, it is indeed possible to differentiate between the empty type-I cohesin on the microbe (OlpA and OlpC) and on the detached cellulosomes (CipA). It is also expected that unoccupied type-I cohesins on CipA would be found close to the CipA-CBM3a and are therefore considered to be colocalized. From our data, it appears that most of the unoccupied type-I cohesins are located on the microbes and should correspond to OlpA and OlpC, given that they are not in the vicinity of CipA-CBM3a, especially on detached cells or in the log phase. It is also interesting to note that the size of the clusters associated with the unoccupied type-I cohesin does not change over the course of growth or when the bacteria are grown on different substrates. OlpA and OlpC have been hypothesized to be involved in shuttling enzymes to cellulosomes [31], but from our data there is no indication that this would be the case. This conclusion is further reinforced, given the fact that the abundance of unoccupied type-I cohesins does not change over time, at least within the constraints of our experiments.

One notable phenomenon that occurs in the stationary phase of growth is the sheer saturation of substrate with cellulosomes. Indeed, the bacterial cells appear to shed cellulosomes over time, and they are clearly being shuttled to the biomass leaving very few unoccupied spots on the surface of the substrate. Again, this phenomenon is consistent with early TEM studies [3, 15] and could potentially explain some of the slowdowns that happen during growth of *C. thermocellum* on insoluble substrates, whereby the saturation of the substrates with cellulosomes could be counterproductive and lead to “traffic jams” of the cellulases on the substrate. Additionally, in all cases, the concentration of cellulosomes, whether on the microbe during log phase or on the substrate during stationary phase, is much higher than anticipated.

This study highlights the potential of using these new optical techniques, combined with specific mutations in *C. thermocellum* or other cellulosomal microbes, to probe and challenge other hypotheses that have been proposed regarding cellulosomes and their mode of action, such as the role of “shuttle” scaffoldins and the role of the attachment of *C. thermocellum* to biomass. These advanced imaging tools will also serve to optimize the process of cellulose deconstruction by these microbes in pure cultures or in cocultures. We note that for cellulases, more is not necessarily better in industrial settings and that controlling cellulosome expression levels could ultimately be important for better biomass deconstruction in these settings.

## Materials and Methods

### Data analysis

Super high-resolution techniques such as PALM and STORM incorporate single-molecule identification and generate large data sets consisting of the molecular coordinates (x-y location) of each molecule detected. Modern-day algorithms exist to identify and locate these molecules but lack the ability to characterize any potential pattern generated by the molecules. Here we utilize unsupervised machine learning (also referred to as cluster analysis) to determine if there are any patterns formed with the distribution of cellulosomes on the bacterial cell wall as well as on cellulose. Unlike supervised machine learning, unsupervised learning does not utilize a training set, but instead finds previously unknown patterns within the dataset without pre-existing labels, and is widely used in clustering, dimension reduction and data representations [33, 34]. Within the field of unsupervised machine learning, there exist multiple techniques ranging from hierarchical clustering, k-means, Density Based Spatial Clustering of Applications with Noise (DBSCAN) and Ordering Points To Identify the Clustering Structures (OPTICS) and others.

Cluster analysis has been utilized within the context of super high-resolution techniques, as described by Sander and coworkers [35], clustering, i.e. grouping the objects of a database (in this case the single molecules) into meaningful subclasses. In this work, we decided to use DBSCAN cluster analysis, developed originally by Ester and coworkers [36]. As defined by these authors, the key idea behind the DBSCAN algorithm is that for each point of a cluster the neighborhood of a given radius has to contain at least a minimum number of points, *i.e.*, the density in the neighborhood has to exceed some threshold [36]. In addition, the shape of the neighborhood is determined by the choice of a distance function for two points [36]. Therefore, DBSCAN only requires two parameters, the radius of a defined area and the MNM within that defined area, which allows the discovery of clusters of arbitrary shape. We utilized DBSCAN from the scikit-learn open-source machine learning library for the Python programming languages [37]. Scikit-learn supports supervised and unsupervised learning and provides various tools for model fitting, data preprocessing, model selection and evaluation, and many other utilities [37].

### Image processing

Multiple software packages were used in the analysis of these datasets, including Matlab for the processing of the PALM/STORM raw data, ImageJ fiji [34], as well as python with the following packages: pandas, opencv, pillow, numpy and matplotlib. Fiji was primarily used to view the processed images generated from the Matlab code as well as the white light images. Fiji was also used to perform image processing, including resizing of the image and merging the different fluorescent images with the white light image. Once the images were processed, they were fed into the python script, written to allow us to select regions of interest (ROI) and capture the ROI from the .csv files, containing the coordinates of each biomolecule. These scripts allowed us to only perform cluster analysis within the ROI, thus saving computational time and reducing the influence of noise. Following the selection of the ROI, the x-y coordinates from each of the biomolecules were then analyzed using python and the scikit-learn-DBSCAN algorithm to characterize their clustering.

### Fluorescent probes

Three fluorescent probes were chosen for the project, based upon the wavelengths available with the Zeiss Elyra super high-resolution imaging. Those chosen were Alexa Fluor 647 [23, 25], and PA-GFP [28]. Alexa Fluor 647 was chosen due to its high blinking efficiency and overall use for STORM imaging. PA-GFP was chosen because regular GFP had been expressed with a dockerin domain that was associated with the type-1 dockerin. Therefore, we employed the mutation by switching the histidine with the tyrosine [28]. This mutation was performed in an *Escherichia coli* expression system, originally designed for the type-1 dockerin [38].

### Primary Antibody expression

Polyclonal antibodies against the CBM3a of the CipA scaffoldin were elicited in rabbits and purified according to the procedure described in Morag et al [39].

### GFP expression

To express the modified GFP, we utilized a plasmid using Q5 High-Fidelity DNA polymerase (New England BioLabs, Ipswich, MA, U.S.A.) according to the manufacturer’s instructions. The codon-optimized CtDoc-GFP expression vector was synthesized by GeneScript (Piscataway, NJ, U.S.A.). pDCYB 209, containing the point mutation (T203H) at amino acid position 203 of the GFP coding sequence as shown in the Supplental document figure S5, was generated using overlapping polymerase chain reaction (PCR) [40]. A 6.514-kb DNA fragment was synthesized with the forward primer DCB619 (ACCTGTCGCATCAATCTGCCCTTTCGAAAGATCCCAACGA) and reverse primer DCB620 (AGGGCAGATTGATGCGACAGGTAATGGTTGTCTGGTAAAAGGAC) using the CtDoc-GFP expression vector as a template. *E. coli* strain DH5a cells were transformed by electroporation in a 2-mm path-length cuvette at 2.5 V, and transformants were selected for kanamycin resistance. The sequences of pDCYB 209 were confirmed by automatic sequencing (Genewiz, NJ, U.S.A.).

*E. coli* BL21(DE3) was used for protein expression. Recombinant strains were grown in LB broth supplemented with kanamycin (25 µg/mL). Cultures were induced at 15°C with 0.2 mM IPTG when OD_600_ = 0.4. Cultures were centrifuged at 5,000 x g for 10 min when OD600 ≥ 1.2. To recover intracellular protein, centrifuged cells were enzymatically lysed in Buffer A (50 mM Tris-HCl, 40 mM NaCl, 10 mM Imidazole, pH 8.0) supplemented with lysozyme, protease inhibitors, and nuclease (Pierce, Waltham, MA, U.S.A.) at 4°C. Enzymatically lysed cells were subjected to 1 min of sonication in a water bath at 10 s intervals punctuated by 30 s on ice. Lysed cells were centrifuged at 10,000 x g for 30 min. The lysate was further purified via immobilized metal affinity chromatography purification using a 5 mL HisTrap FF Crude column (GE Healthcare, Piscataway, NJ, U.S.A.). Once bound, Buffer B (Buffer A with 200 mM Imidazole) was used to elute the protein. The recovered fractions were then purified further via size exclusion chromatography using a HiLoad 16/600 Superdex 200 prep grade column (GE Healthcare, Piscataway, NJ, U.S.A.). This step served to buffer exchange the fractions into SEC Buffer (200 mM acetate, 100 mM NaCl, 10 mM CaCl_2_, pH 5.5). Once collected, the recovered protein was concentrated using a 10-kDa spin concentrator (Sartorius, Stonehouse, UK). Final protein concentration was determined using a Pierce BCA protein assay (Pierce, Rockford, IL). Purified CtDoc-GFP(PA) protein was analyzed by SDS-PAGE using a 4-12% NuPAGE Bis-Tris Gel (Invitrogen, Carlsbad, CA, U.S.A.) run at 150 V for 55 min in MOPS SDS buffer. The gel was then stained with Colloidal Blue (Invitrogen, Carlsbad, CA, U.S.A.).

### Bacterial growth and characterization

Growth studies were performed with DSM1313 and CTN5 strains in 50 mL total volume in anaerobic serum bottles, containing 20 mL of CTFUD media with 2% D(+)-cellobiose (ACROS Organics) or 0.5% Avicel PH-101 (Sigma-Aldrich) as the carbon sources. Seed cultures were grown at 60°C from frozen stock in 20 mL of culture medium, containing 0.5% cellobiose CTFUD media. Upon reaching exponential phase, samples were transferred from seed bottles to fresh bottles containing either 2% cellobiose or 0.5% Avicel, where the initial optical density (OD) was set at ∼0.045. Transferred cultures were grown at 52°C with agitation (∼100 rpm for cellobiose bottles, ∼200 rpm for Avicel bottles). The cultures were grown overnight for 5 to 8 h, after which samples were taken periodically until the cultures reached steady state.

OD sampling was performed using a Genesys 20 spectrophotometer (Thermo Scientific) at 600 nm. When sampling Avicel bottles, 1.5-mL samples were centrifuged at 0.3 min^-1^ x g for 30 s to precipitate the Avicel. The supernatant fluids were sampled using a pipette and used to determine the sample OD.

### Chromophore and GFP attachment

For the two chromophores, two primary antibodies were expressed, one primary antibody expressed in rabbit for the CBM3a located on the CipA scaffoldin and another primary antibody expressed in rabbit that is associated with the GH48 enzyme. Wildtype or the mutant bacterial cells were grown with several passages during log phase to ensure a healthy culture. Once the bacterial cells were in the appropriate growth stage (either log phase or stationary phase) 2 mL of the bacterial cultures were extracted from the serum bottles, centrifuged and washed three times using 50 mM SEC buffer supplemented with 5 mM CaCl. The cells were then resuspended in the SEC buffer, both primary antibodies were added, and the suspension was incubated at room temperature for 5 min. The immunolabeled bacterial cells were washed an additional three times with the SEC buffer by centrifugation as described above. The primary antibody-tagged bacterial cells were then incubated with the secondary antibody tagged with AF 647 along with the PA-GFP for 5 min. The bacterial cells were washed by centrifugation a final three times in PBS buffer and stored in PBS buffer to match the imaging buffer used in the experiments. Sample preparation and imaging occurred on the same day. Labeled bacterial cells were drop cast onto a poly-L-lysine-coated 25-mm round glass coverslips.

## Acknowledgment

This material is based upon work supported by the Center for Bioenergy Innovation (CBI), U.S. Department of Energy, Office of Science, Biological and Environmental Research Program under Award Number ERKP886. This work was authored in part by Alliance for Sustainable Energy, LLC, the Manager and Operator of the National Renewable Energy Laboratory for the U.S. Department of Energy (DOE) under Contract No. DE-AC36-08GO28308. The views expressed in the article do not necessarily represent the views of the DOE or the U.S. Government. The U.S. Government retains and the publisher, by accepting the article for publication, acknowledges that the U.S. Government retains a nonexclusive, paid-up, irrevocable, worldwide license to publish or reproduce the published form of this work, or allow others to do so, for U.S. Government purposes.

## Spatial resolution of the super-high resolution microscope

For conventional microscopy, the lateral resolution is given calculated by the Abbe Equation [1]

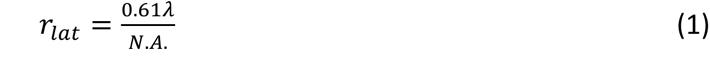

And the axial resolution is defined as [1]

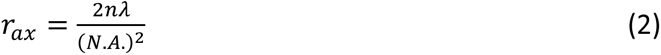

Super-high-resolution techniques enable microscopy to break the diffraction limit by temporally or spatially modulating the excitation or activation of light via the chromophores. Data collected from the super high resolution microscope were generated by iterating the optical images and capturing individual cells in both the log phase and the stationary phase of growth. Each bacterial cell was analyzed with the DBSCAN clustering algorithm by keeping the epsilon constant at a 75-nm radius and iterated through the MNM. Because the various bacterial cells did not necessarily have the same number of clusters, nor did they have the same MNM, the data from the DBSCAN cluster analysis for each bacterium were normalized, and the normalized data were averaged among all other bacterial cells.

Figure S1 shows a normal fluorescent image of bacteria attached to Avicel (S1A). The bacteria are labeled with the AF 647 chromophore attached via an CBM3 antibody located on the cellulosome. The fluorescence is diffuse resulting in the difficulty in distinguishing its structure as compared the super-high-resolution image in figure 1B where fluorescent events can be distinguished resulting in a sharper image. The purple arrow highlights the region used in figure 1C demonstrating the true optical spatial resolution of the STORM/PALM super high-resolution microscope used in this work. There are two circled regions (region C1 and region C2) with the actual measurements shown. Region C1 shows 4 distinct fluorescent chromophores within a total distance of ∼130nm.

**Figure S1.**
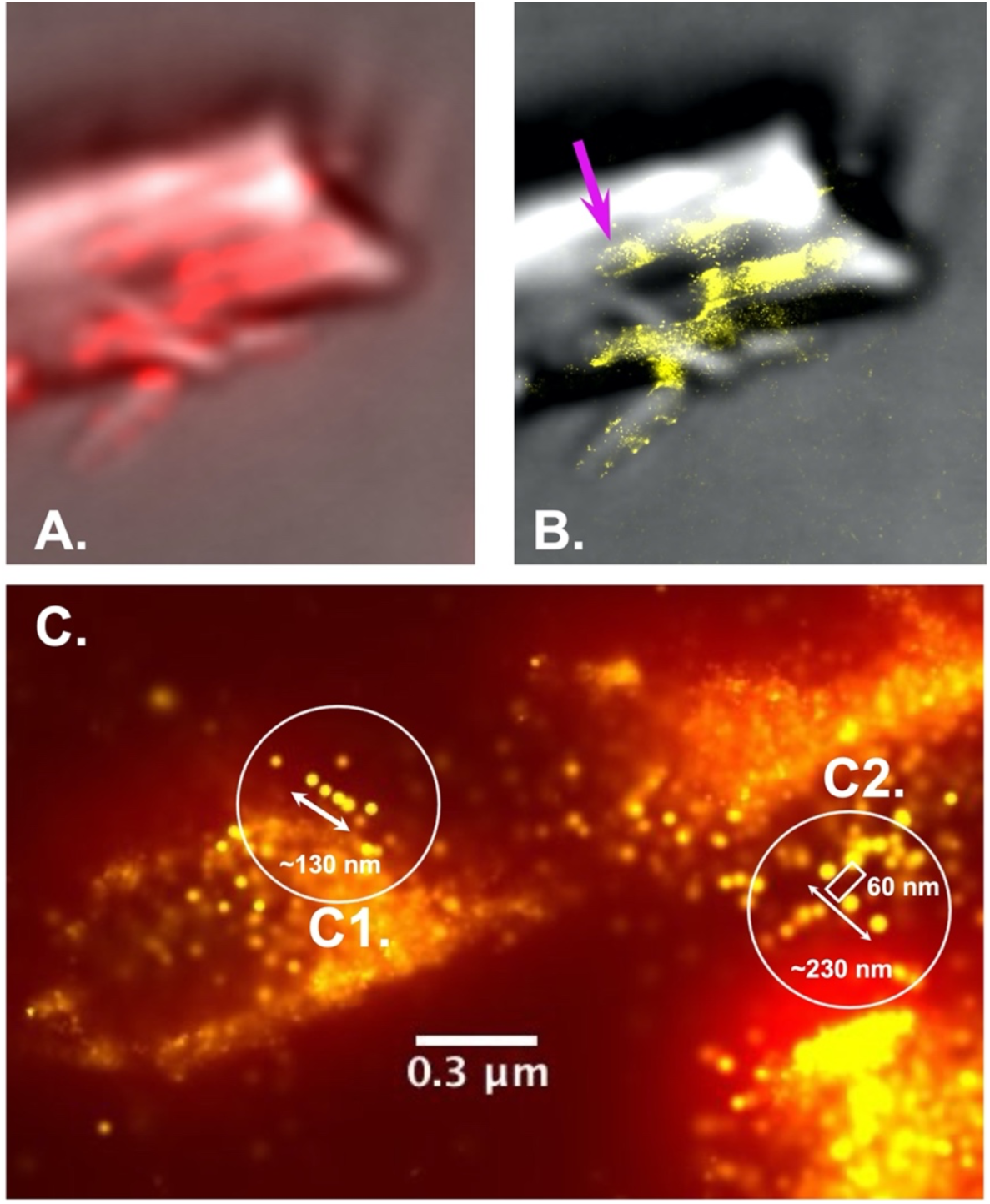
Super high special resolution for the STORM microscope using an Alexa Fluor 647 chromophore attached to a secondary antibody associated with the CBM3 located on the primary scaffold. Figure 1A is the far field fluorescence associated with microscopy with low resolution. Figure 1B is the fluorescent signal from the super high resolution STORM image where higher spatial resolution can be achieved in comparison to the low-resolution fluorescence image in A. Figure 1C demonstrates the actual measured spatial resolution with multiple fluorescent events observed within 100 to 200 nm.

This highlighted region demonstrates the ability of the system to image below the diffraction barrier of light. These 4 fluorescent events within a low-resolution system would appear as one larger unresolved gaussian fluorescence signal. Region C2 shows the measured spacing of the three fluorescence events to be 60nm, well below the diffraction barrier of light.

Similar spatial resolution is demonstrated with the photoactivated GFP (PA-GFP) as shown in figure S2. Figure 2a is the white light image in conjunction with the fluorescence from the PA-GFP. Figure 2b is the fluorescence from the super high-resolution microscope with the crisp single fluorescent events. Figure 2C shows a selected region of a single bacterium with the PA-GFP attached to a vacant type-1 dockerin site. The region in the circle shows three PA-GPFs separated by ∼100nm with a distance end to end of ∼280nm. This helps demonstrate that the spatial resolution of both the STORM system (AF647) and the PALM system (PA-GFP) is below the diffraction barrier of light.

**Figure S2.**
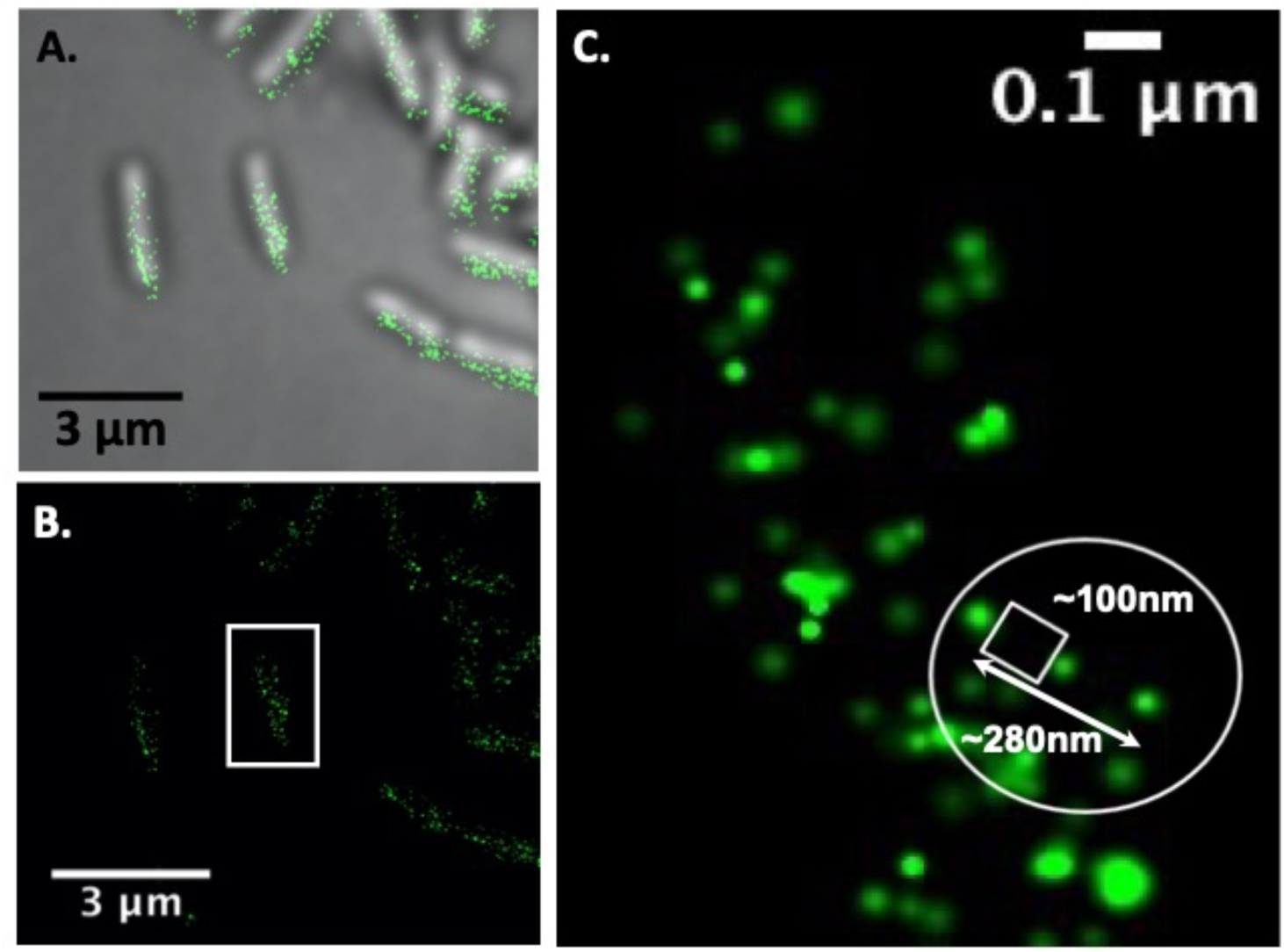
Super high resolution for PALM microscopy using the PA-GFP for the Protein-dockerin fusion protein (PA-GFP-Dockerin) targeted to the OlpC site on the bacterial cell wall as well as on the cellulosome. Figure S2A shows the far field fluorescence associated with the low resolution microscope image. Figure S2B is the fluorescent signal from the super high resolution PALM imaged where the special resolution can be achieved in comparison to the low-resolution fluorescence image in S2A. Finally, S2C demonstrates the actual measured spatial resolution within of less than 100 nm

## Quantitative vs Qualitative microscopy imaging

Traditional optical microscopy does not possess the ability to systematically quantify information within the images and the interpretation of the optical microscopy images is more subjective. Figure S3 demonstrates the visual difference of the fluorescence pattern between *C. thermocellum* in log phase (figure S3a) and in stationary phase (figure 3b.) Figure 3 shows 18 individually selected bacterial sells from the over 60 bacterial cells used in the DBSCAN clustering analysis. Visually, there is a difference between the log phase bacterial cells and the stationary bacterial cells. The issue arises in being able to describe the overall distribution pattern of the fluorescence on the surface of the bacterial cell wall. Visually speaking, the log phase bacterial cells (figure 3a) appear to have an increase in fluorescence on the bacterial wall compared to bacterial cells in stationary phase (figure S3B). From this perspective, we can only speculate if there is truly a reduction in the fluorescence and from figure 3B, out of 18 bacterial cells, 60% appear to have reduced fluorescence. Using our approach, we can perform quantitative analysis on everything single bacterial cell and develop a distribution pattern of the total number of cellulosomes on the surface of the bacteria and correlate with the different phases of growth.

**Figure S3.**
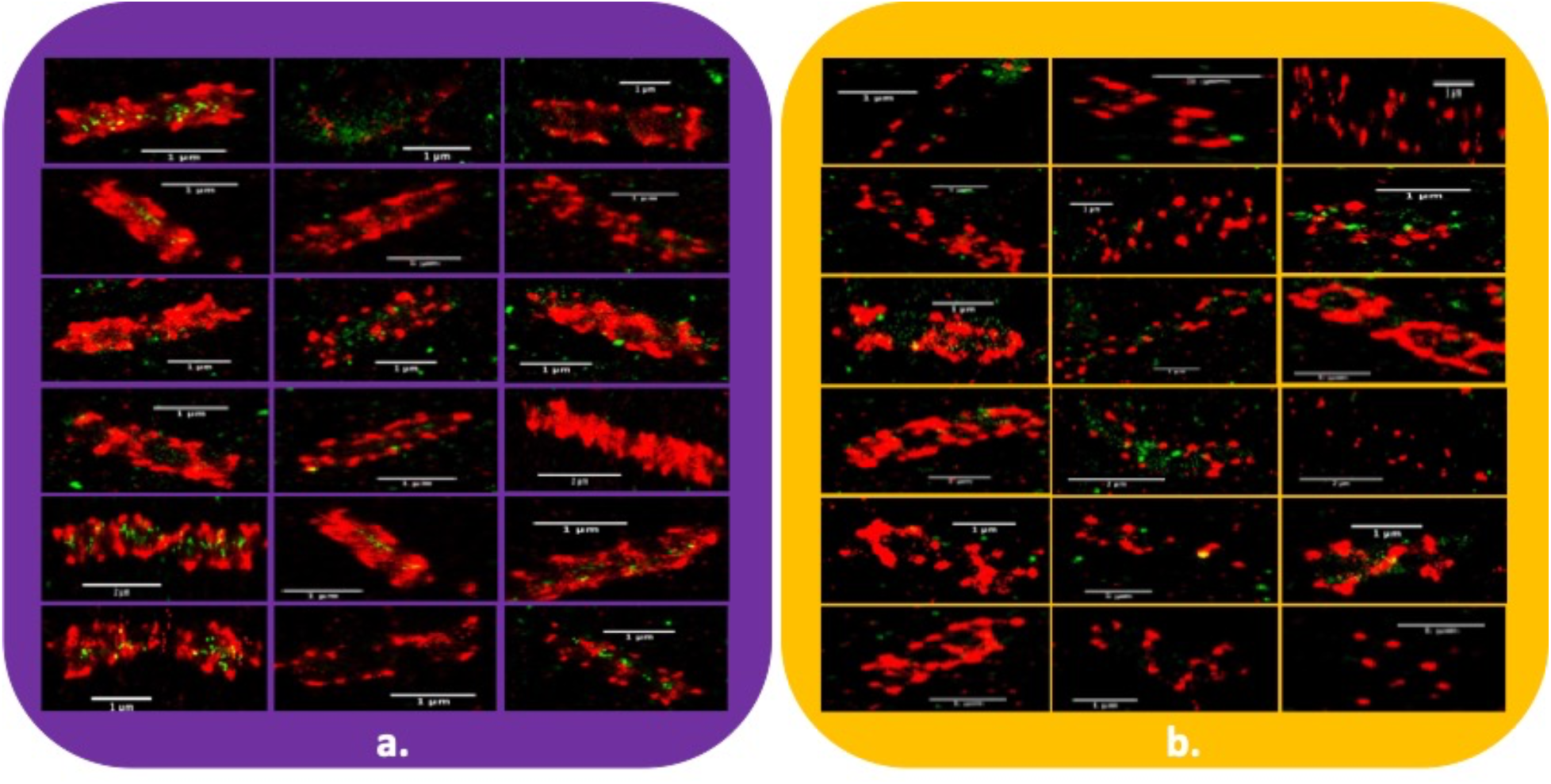
Qualitative representation of single C. thermocellum bacterial cells in log phase (a) and stationary phase (b). Here, we have chosen to show 18 selected images of bacterial cells during in log phase and stationary phase. There does appear to be a difference in the pattern of fluorescence between the two growth stages, but quantitative imaging is required to draw these conclusions.

Figure S4 also demonstrate the same issue when looking at bacterial cells on a substrate. Figure S4a shows the fluorescence pattern of cellulosomes on both the bacteria and the Avicel during log phase and figure 4b shows the fluorescence of the cellulosomes on both the bacteria and Avicel during stationary phase. Once again, these are selected images taken from the two datasets. The complex nature of the fluorescence distribution during log phase demonstrates the need to implement the DBSCAN analysis as it is difficult to distinguish the boundary between the Avicel and the bacterial cells.

**Figure S4.**
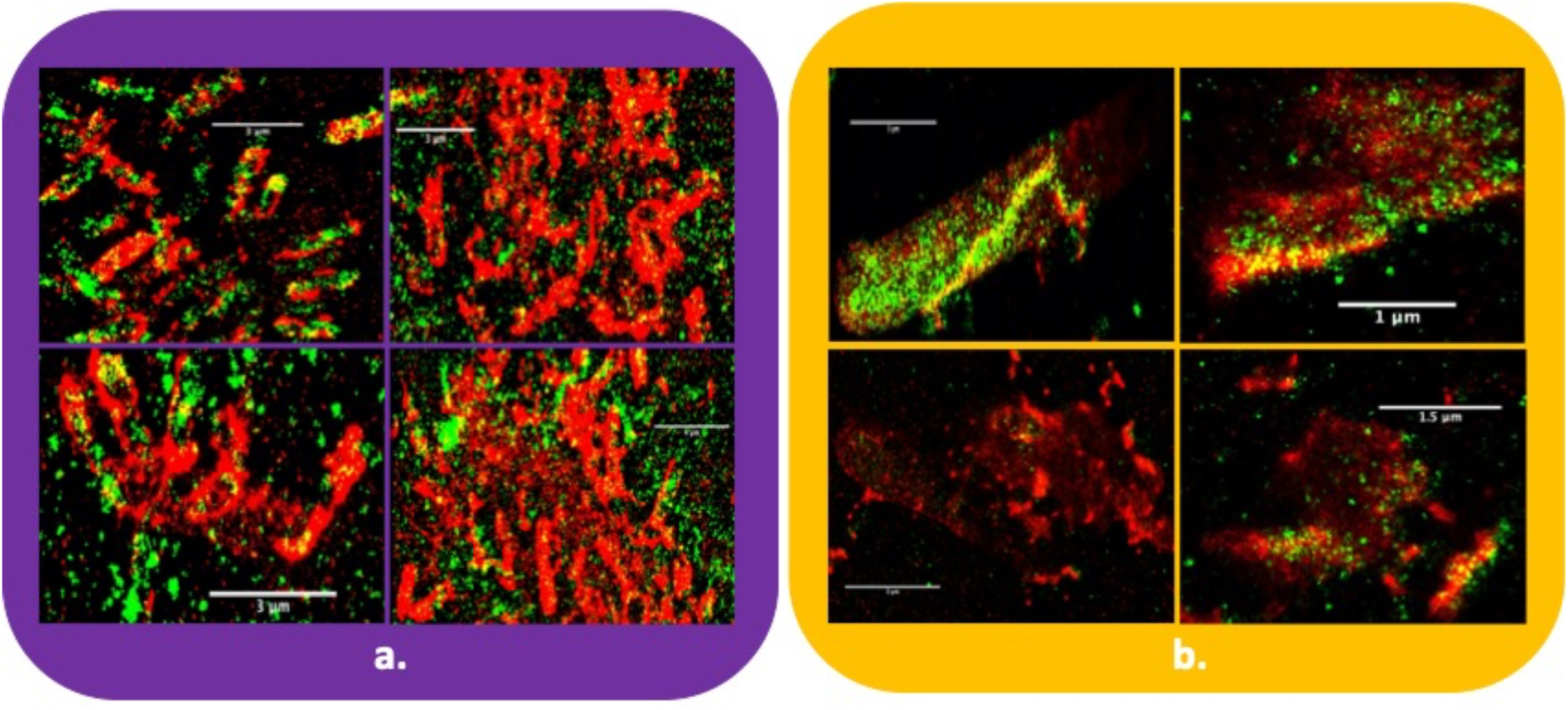
Qualitative representation of C. thermocellum bacteria cells on Avicel in log phase (a) and stationary phase (b). Here, we have chosen to show 4 selected images of bacteria cells during log phase and stationary phase. 4a demonstrate the complex nature of the fluorescence distribution pattern on both Avicel and the bacterial cells, making it rather difficult to describe what the fluorescence distribution pattern looks like.

## PA-GFP Expression for super high resolution PALM imaging

The Photoactivated Green fluorescent protein (PA-GFP_ was express a plasmid using Q5 High-Fidelity DNA polymerase (New England BioLabs, Ipswich, MA, U.S.A. The codon-optimized CtDoc-GFP expression vector was synthesized by GeneScript (Piscataway, NJ, U.S.A.). pDCYB 209, containing the point mutation (T203H) at amino acid position 203 of the GFP coding sequence as shown in the figure S5, was generated using overlapping polymerase chain reaction (PCR) [2]. *E. coli* strain DH5a was used as the transformant.

**Figure S5.**
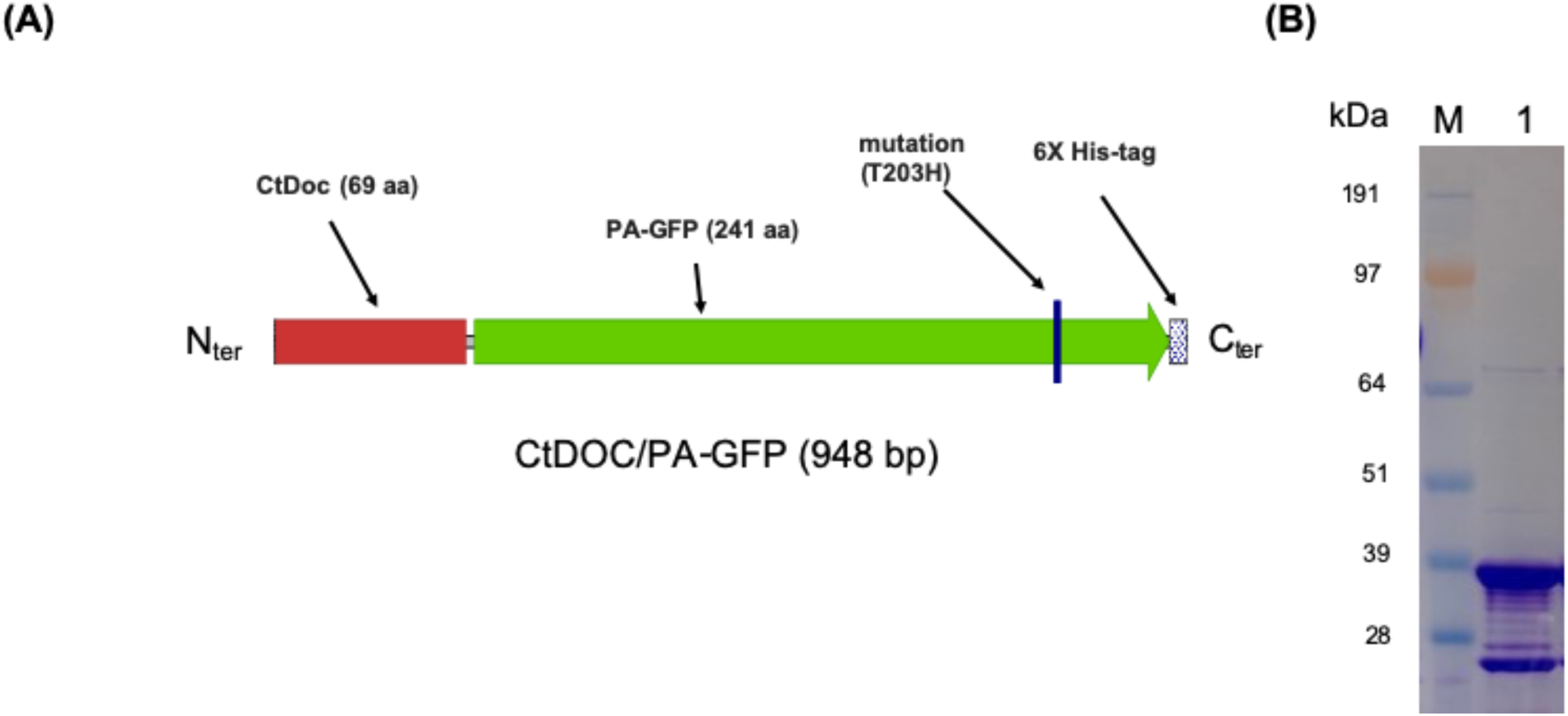
Diagrammatic representation of the CtDoc-GFP(PA) construct and confirmation of its expression. (A) The size and location of the CtDoc (Type-I dockerin domain derived from the C. thermocellum cellulosome), PA-GFP (Photoactivated GFP), the Thr to His point mutation, and the 6X Histag. All features are drawn to scale. (B) Expression and purification of CtDoc/PA-GFP (̴37 kDa) in E. coli. M: Invitrogen SeeBlue Plus2 Prestained Standard; lane 1: Purified CtDoc/PA-GFP.

